# Soil causes gut microbiota to flourish and total serum IgE levels to decrease in mice

**DOI:** 10.1101/2021.01.25.428035

**Authors:** Dongrui Zhou, Na Li, Fan Yang, Honglin Zhang, Zhimao Bai, Yangyang Dong, Mengjie Li, Wenyong Zhu, Zhongjie Fei, Pengfeng Xiao, Xiao Sun, Zuhong Lu

**Author notes:** **For Correspondence**: Key Laboratory of Child Development and Learning Science of Ministry of Education, Southeast University, Nanjing 210096, China.

## Abstract

Traditional farm environments induce protection from allergic diseases. In this study, farm environmental factors were classified into three categories, environmental microbes, soil, and organic matter. To explore the impact of soil and environmental microorganisms on gut microbiota and immune function, mice were fed sterilized soil, soil microbes (in lieu of environmental microbes), or non-sterilized soil. Metagenomic sequencing results showed the intake of sterile soil i.e. inhaling a small amount of soil microbes in the air increased gut microbial diversity and the abundance of type III secretion system (T3SS) genes, and decreased serum immune globulin E (IgE) levels induced by 2-4-dinitrofluorobenzene(DNFB). The intake of soil microbes increased the abundance of genes involved in the metabolism of short chain fatty acids and amino acid biosynthesis. Meanwhile, the intake of soil increased gut microbial diversity, the abundance of T3SS genes and related infectious elements, and genes associated with the metabolism of short chain fatty acids and amino acid biosynthesis, and decreased serum IgE levels. Therefore, soil may be useful as a potential “prebiotic” promoting the reproduction and growth of some intestinal microorganisms that harbor bacterial secretion system genes, especially those of T3SS, whose abundance was positively and significantly correlated with innate immune function of mice.

## Introduction

Several epidemiological studies have shown that children growing up on traditional farms suffer less from asthma, hay fever, and allergic sensitization (Riedler et al., 2001; Mutius and Vercelli, 2010; Ege et al., 2011; Lehtimäki et al., 2020). A healthy gut microbiota is necessary for proper human immune function (Schuijs et al., 2015; Honda and Littman, 2016; Vatanen et al., 2016; Stephen-Victor and Chatila, 2019). Many studies suggest that farm environments could increase the diversity and richness of gut microbiota and shape its composition or structure (Schuijs et al., 2015; Zhou et al., 2015; Vatanen et al., 2016; Gilbert and Blaser, 2018; Noora et al., 2018), especially that of children who live on farms at an early age (Adlerberth and Wold, 2009; Wopereis et al., 2014).

There is a high-level of exposure to airborne microbes, animals, dust, plants, and soil on farms (Rook, 2013). In addition, a wide variety of different microorganisms are present throughout farms (Rook, 2013; Mhuireach et al., 2016). It has been speculated that the environmental microbiota present in ambient air may interact with and supplement an individual’s intestinal microbiota (Rook, 2013). Farm animals are also considered an important factor that may contribute to improving human immunity (Debarry et al., 2007; Vogel et al., 2008) and the microbiota in dust of households with pets is substantially richer and more diverse than found in homes without pets (Fujimura et al., 2010). Furthermore, cohabiting individuals tend to share gut microbial communities (Song et al., 2013), but interestingly, people tend to share more microbial communities with their dogs (Song et al., 2013). It has been reported that the field area of green space is inversely proportional to the incidence rate of allergic diseases (von Hertzen et al., 2011; Kembel et al., 2012). This may be explained by plants shaping the content and abundance of rhizosphere microorganisms and provide habitats for the microorganisms on the ground (Berendsen et al., 2012). Finally, there is a lot of dust on farms with the principal components being microbes, soil, pollen, animal dander, and hair (Rook, 2013).

As an important component of the farm environment, soil is the microecological environment with the largest number of microbial species on earth (Blum et al., 2019) and is known as the “seed bank” of microorganisms(Lennon and Jones, 2011). In comparison, the number of different species of human intestinal microorganisms is approximately 1/10^th^ found in soil (Blum et al., 2019). Interestingly, approximately 80% of the microorganisms in the soil are in a dormant state, similar to plant seeds (Blum et al., 2019). Once they are exposed to a suitable living environment, they multiply (Lennon and Jones, 2011). In primitive farming stages and less developed areas, people have greater contact with soil (Zhou et al., 2018), children often use their mouths to explore their surrounding environment (A, 2003; Zhou et al., 2018), and in some countries, individuals actually use soil as a food source (Sing and Sing, 2010). Finally, soil mixed in the bedding material or present in the living environment of animals has an important effect on their gut microbiota (Zhou et al., 2018; Grieneisen et al., 2019).

To investigate the specific factors on farms that influence gut microbiota, we grouped farm environmental factors into three categories, environmental microbes, soil, and organic substances from plants or animals (Table 1). To explore the impact on the gut microbiota of each factor, excluding organic matter, adult mice (Table S1) were randomly divided into four groups, a group fed soil (Soil), a group fed sterilized soil (SS), a group administered with microbes in their water (MW), and a normal control group. The animals were analyzed using 16S rDNA and metagenomic sequencing to evaluate the composition and richness of the gut microbiota. The specific treatment of each group is shown in Fig. 1A and Fig. S1. To test the effect of the treatments on immune function, 2-4-dinitrofluorobenzene (DNFB) was used topically to induce eczema on the skin of the mice, and the serum IgE levels were then measured.

**Table 1.**
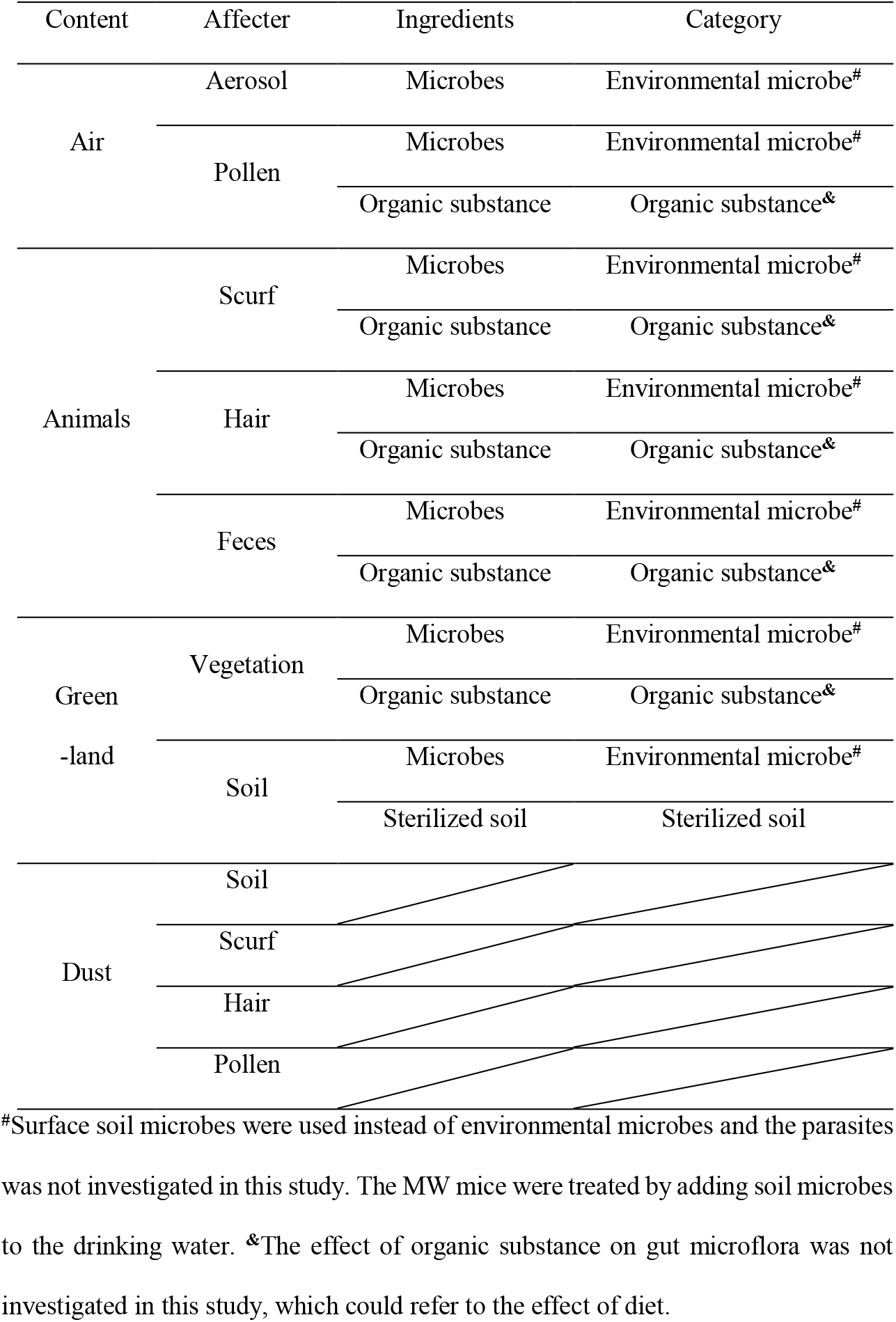
Summary of farm environmental factors affecting gut

**Fig. 1.**
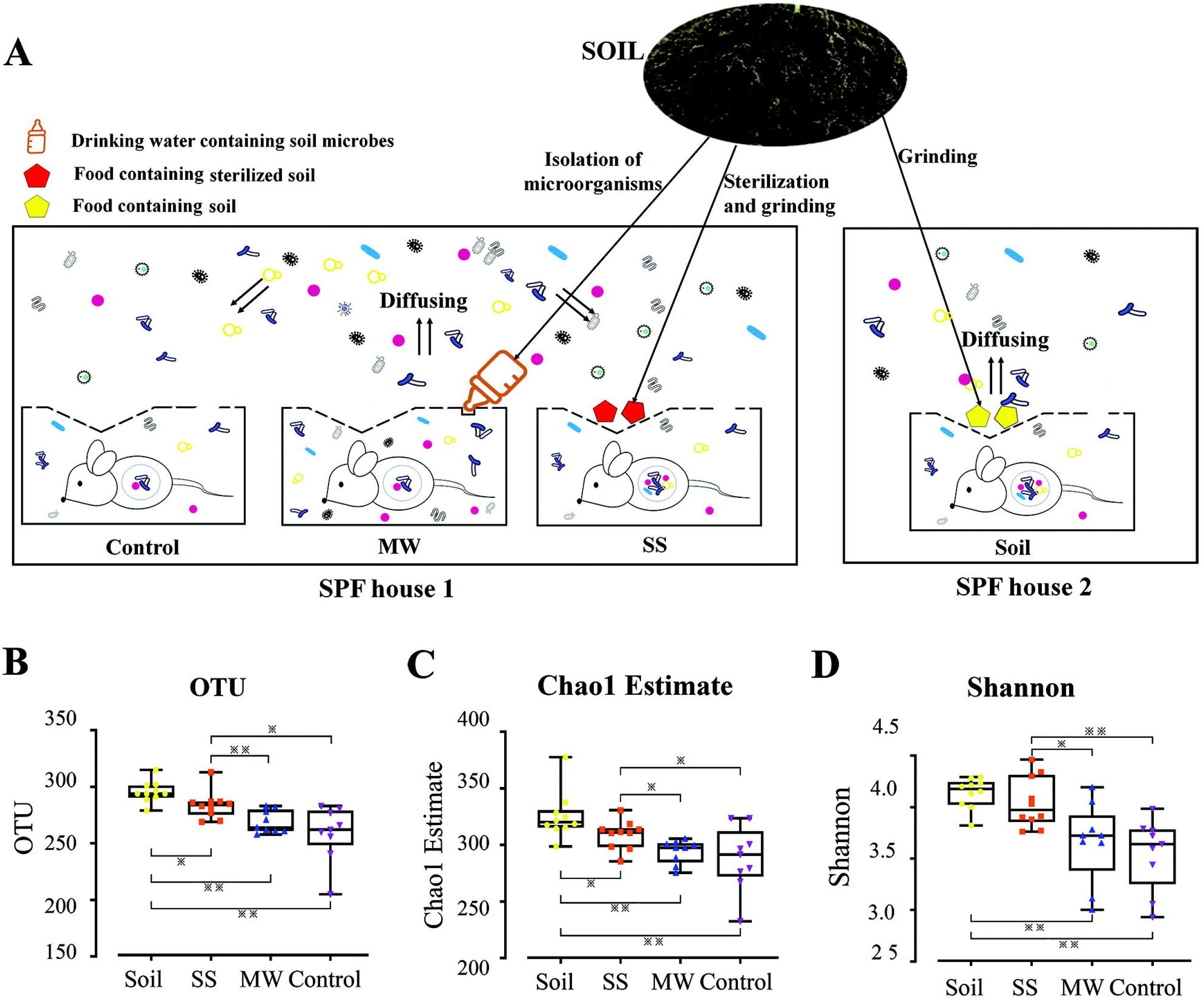
Schematic diagram of mouse groups and experimental treatments, and Richness and diversity of the four experimental groups of mice. A. Control: experiment control mice; MW: mice provided drinking water containing soil microbes (Table S2); SS: mice fed diets containing sterilized soil (Table S3); Soil: mice fed diets containing unsterilized soil. The Control, MW, and SS groups of mice were housed in the same SPF room. All mouse cages were covered with wire mesh, which allowed microbes to diffused from the MW mouse cages. The Control and SS mice inhaled microbes from the cages of the MW mice. The Soil group of mice were housed in different SPF animal room. Diet, age, nest, lighting, cleanliness, bedding, house temperature, and humidity were consistent for all four groups of mice. B. Box graph of unique operational taxonomic units (OTUs) at a 97% threshold. C. Chao1 estimators. D. Shannon index. (n = 9 or 10; \**P* < 0.05, \*\**P* < 0.01, two-tailed least significant difference test)

## Results

### Soil-intake or sterilized-soil-intake increased gut microbial diversity and richness

The four groups of mice were treated for 42 days as shown in Fig. 1A. Fresh fecal samples from mice were collected (Fig. S1) and the gut microbiota analyzed using high-throughput sequencing.

We sequenced the 16S rDNA of the fecal samples with polymerase chain reaction (PCR) amplification of the V4 hypervariable region primed using the 515F-907R primer set. For every quarter increase in microbial diversity or richness, the risk of allergic disease is reduced by 55% (Azad et al., 2015). A total of 1,921,486 sequences were qualified with the average sequence of each sample being 50,565. We used rarefaction to normalize the number of reads in each sample of the operational taxonomic unit (OTU) table to 38,680 sequences and analyzed the diversity of the intestinal microbiota in each group.

As shown in Fig. 1B and Table S4, the number of OTUs in the SS group was significantly more than that in the MW or Control groups (*P* < 0.05), but less than that in the Soil group (*P* < 0.05). There was no significant difference between the MW and Control groups. Similar results were obtained when abundance was estimated using the Chao1 index (Table S4 and Fig. 1C). However, the Shannon index showed no significant difference between the SS and Soil groups (Table S4 and Fig. 1D).

From this viewpoint, the intake of sterilized soil significantly improved the diversity and richness of intestinal microorganisms in mice to levels similar to that of the Soil group. However, neither the diversity nor richness of the gut microbiota significantly changed when only microbes isolated from soil were ingested.

### Changes in gut microbial composition

Principal coordinates analysis (PCoA) of the unweighted UniFrac distance matrix showed obvious differences between each group of mice. Specifically, the intestinal microbial structure of the mice in the SS and Soil groups were similar, whereas those in the MW and Control groups were similar (Fig. 2A). It could also be seen that the distance between the SS and Soil groups was shortest, followed by that between the Control and MW group (Fig. 2B, Table S5). The longest distance was between the MW and Soil groups. Furthermore, PCoA of the Bray-Curtis matrix of metagenomics sequences were like that of the 16S rRNA gene sequences (Fig. 2C and 2D, and Table S5).

**Fig. 2.**
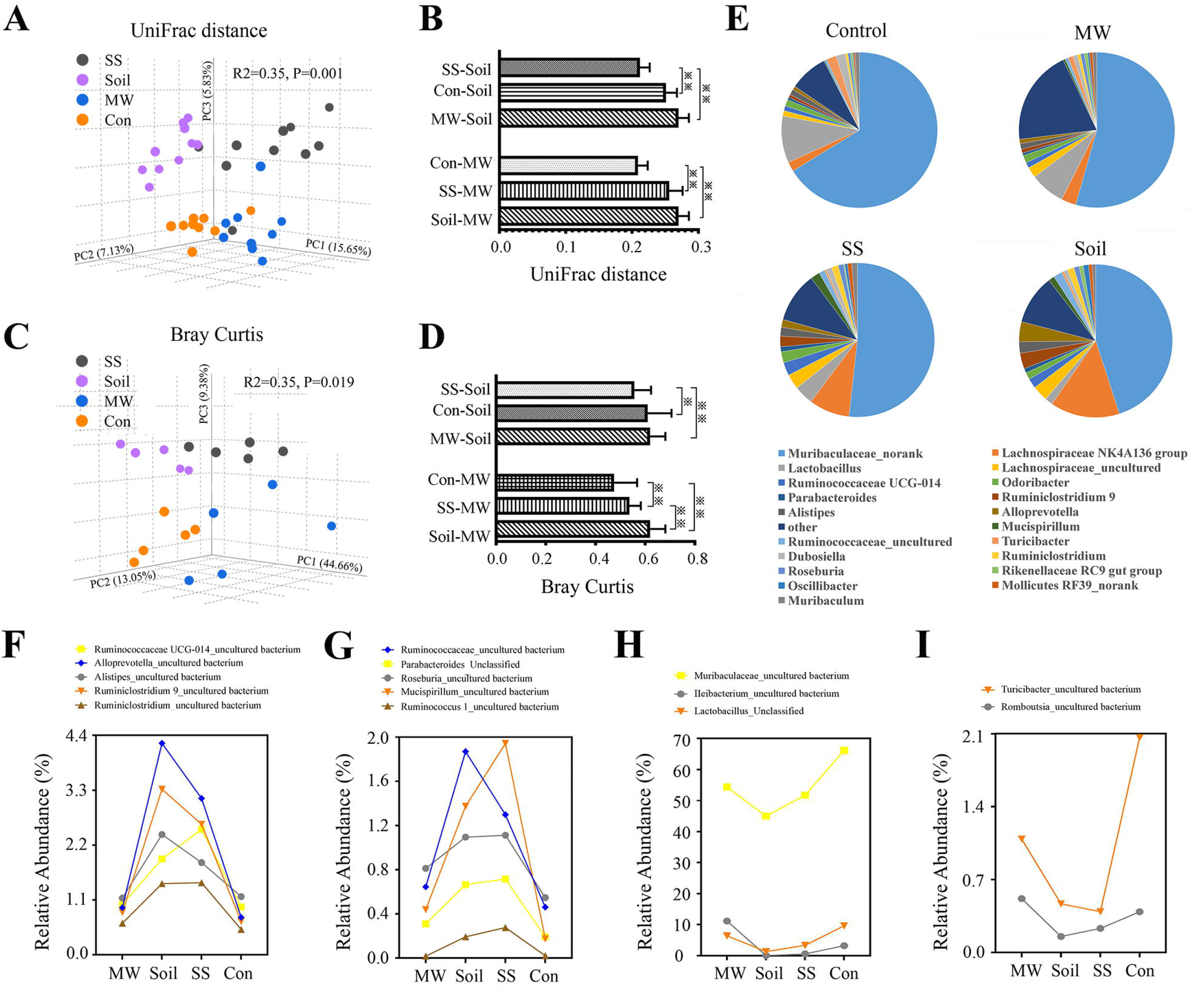
Changes of gut microbial community. A. principal coordinates analysis (PCoA) of unweighted UniFrac distance of 16S rRNA gene sequencing. B. Each bar represents the mean ± SEM (n = 9 or 10). C. PCoA of unweighted Bray-Curtis matrix of metagenomics shotgun sequences. D. Each bar represents the mean ± SEM (n = 5). E. Pie chart of top 20 most abundant genera. \**P* < 0.05, \*\**P* < 0.01, according to two-tailed least significant difference test. F,G,H,I. Species with obvious differences in relative abundance classified via 16S rDNA sequences of the gut microbiota. Strains abundant in the Soil and SS groups of mice (F, G) and the MW and Control groups of mice (H, I). Control: experiment control mice; MW: mice provided drinking water containing soil microbes; SS: mice fed diets containing sterilized soil; Soil: mice fed diets containing unsterilized soil.

The most abundant genera based on 16S rDNA sequences were identified using the RDP Classifier. There were obvious differences in the composition of the top 20 most abundant genera among the four groups (Fig. 2E and Table S6). The pie chart of the Control group was more similar to that of the MW group, whereas that of the SS group was more similar to that of the Soil group. The same conclusions were reached using the column of genera abundance/type in each phylum of the top five most abundant phyla (Fig. S2) and the results of species abundance/type among the different groups (Fig. 2F,2G,2H and 2I).

Random forest is a supervised machine learning technique that uses multiple decision trees to train and predict samples. It is a powerful classifier that can use non-linear relationships and complex dependence between OTUs/strains to identify the OTUs/strains that are important to the structural makeup of the microbiota. An importance score is assigned to each OTU/strain based on the increased error caused by deleting that OTU/strain from the prediction set. In the current study, we considered an importance score for an OTU of at least 0.0005 as being highly predictive. There were 67 predictive OTUs at the species-level between the SS and SPF groups, of which 55 (82%) were overrepresented in the SS group (Table S7). However, there were 36 predictive OTUs between the MW and SPF groups with only 25 OTUs (69%) being overrepresented in the MW group (Table S7). Correspondingly, analysis of the Soil and SPF groups showed there were 74 predictive OTUs with the Soil group presenting 63 (85%) more OTUs (Table S7). Interestingly, compared with MW group, there were 64 and 49 predictive OTUs in Soil and SS groups, respectively, among which 54 (84%) were overrepresented in Soil group and 41 overrepresented in the SS group (Table S7).

The results of random forest analysis of strains based on shotgun sequencing of the microbial metagenome were similar to those of the above 16S rDNA sequencing analysis (Fig. 3 and Table S8). The strains listed in Table S8 meet two standards; first, the random forest importance score of the strain was at least 0.001, and second, the p-value of t-test was less than 0.05. Compared with Control mice, there were 128 predictive species for the Soil mice, 108 for the SS mice, and 52 for the MW mice, overrepresenting 96, 85, and 47 microbes, respectively (Fig. 3A and Table S8).

**Fig. 3.**
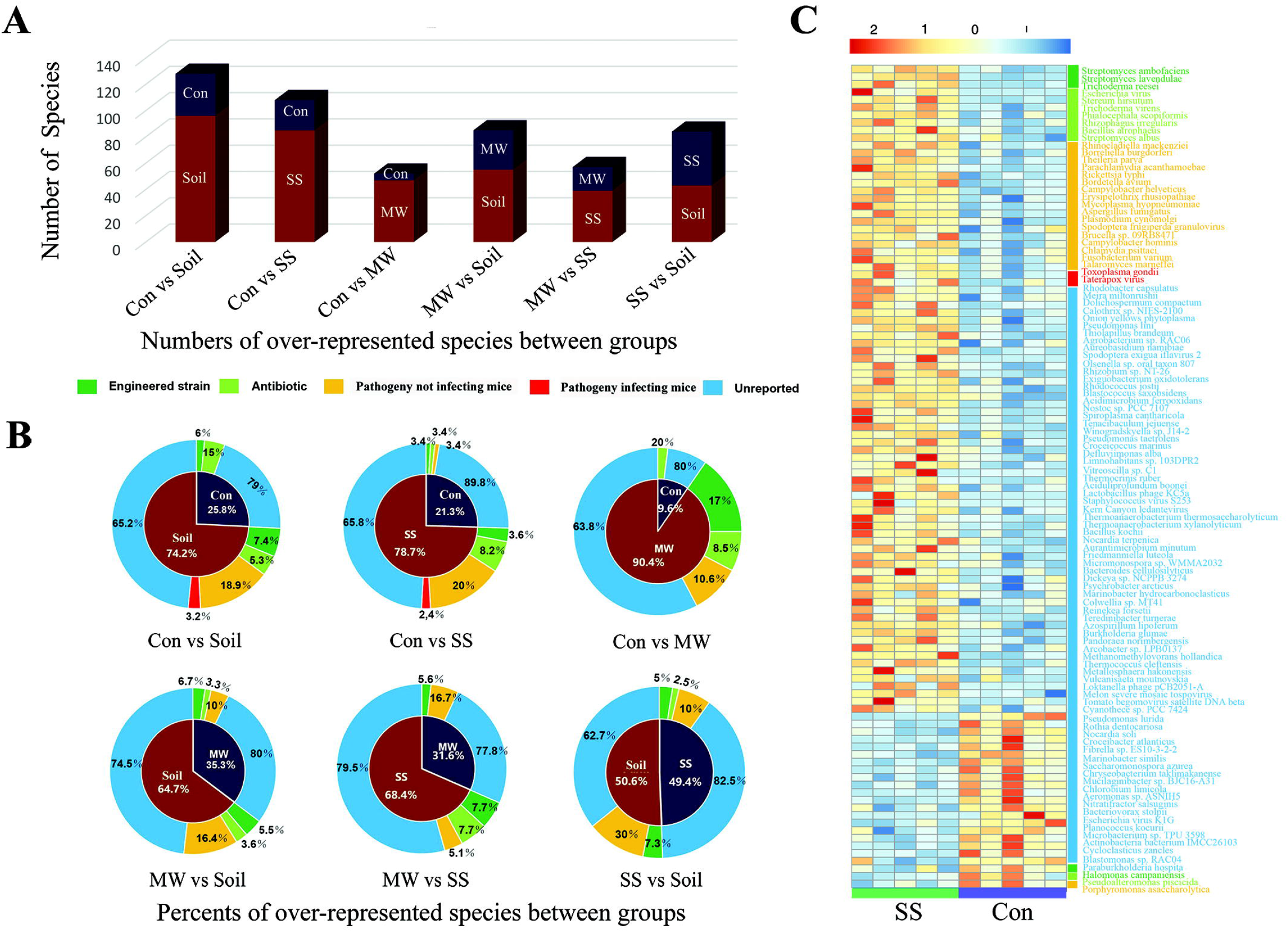
Species identified via random forests analysis based on shotgun sequencing that exhibit significant differences in their representation in the fecal microbiomes of between each two groups of mice. A. Column diagram of the number of over-represented species. B. Pie charts of the ratio of over-represented species and the percent of each functional microbe. C. Heatmaps for impact of sterilized soil-intake on species of gut microbial metagenomics. Each color characteristic represents the same functional microbial species (n = 5/group). The data are listed in Table S8.

Based on these results, we concluded that ingestion of soil-isolated microbes, sterilized soil, or farm soil each had an influence on the intestinal microbial structure and composition of mice. The effect of eating sterilized soil was more similar to that of the Soil group, whereas the effect of drinking soil microbes was more similar to that of the Control group.

### Intake of sterilized soil increased the abundance of type III secretion system (T3SS) genes

To further understand the mechanism by which ingesting soil or drinking soil microbes influenced the mouse intestinal microbiota, analyses of microbe species and genes were conducted using the metagenomic shotgun sequence data. To search for their biological relevance, we selected microbe species with significant differences between experimental groups based on t-test analysis and with importance scores exceeding 0.001 (Fig. 3A, Table S8). It was determined that the biological functions for 62.7–89.8% of the microbes selected had not been previously published (Fig. 3B, Table S8). Among the species selected, four had functions reported. One is an engineering bacterium, the second is an antibiotic producing microbe, the third is pathogenic to plants or animals other than mice, and the fourth one is a mouse pathogen (Fig. 3B,3C and Table S8). Compared to that of the Control mice, the SS and Soil mice intestinal microbiota were more abundant for mouse pathogens (Fig. 3 and Table S8), whereas the MW mice had no greater number of mouse pathogens but had more of four other types of microbes (Fig. 3C and Table S8).

When the abundance of functional genes was compared between groups, it was found that the SS group showed a significant increase in the abundance of genes encoding T3SS and two-component systems compared with that in the Control or MW mice (Fig. 4 and Table S9). Compared to the MW mice, the Soil mice harbored more genes encoding T3SS (Fig. 4 and Table S9), two-component systems, and butanoate metabolism (Table S9). Accordingly, the ingestion of soil or sterilized soil increased the abundance of mouse and other pathogens, as well as genes coding for T3SS and two-component systems.

**Fig. 4.**
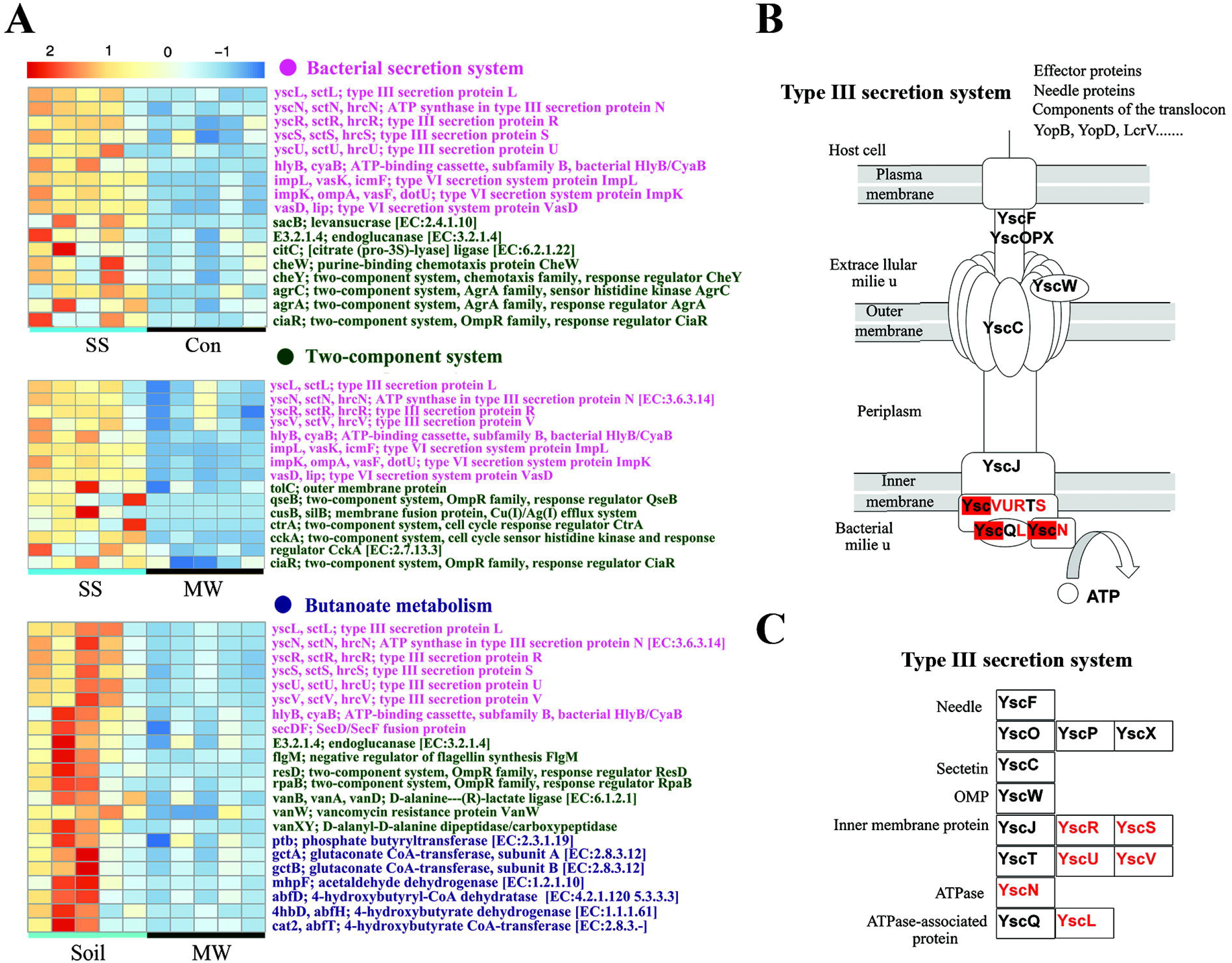
Effects of sterilized soil-intake on gene abundance of gut microbial metagenomics. A. Heatmaps for comparison between the SS and Control groups of mice, SS and MW, and Soil and MW (n = 5/group). B,C. Diagram of Kyoto Encyclopedia of Genes and Genomes (KEGG) entries for the type III bacterial secretion system. KEGG entries whose proportional representation was higher in the fecal microbiomes of the SS and Soil mice compared with that of the Control or MW mice. P-values for the highlighted Kos can be found in Table S9. Control: experiment control mice; MW: mice provided drinking water containing soil microbes; SS: mice fed diets containing sterilized soil; Soil: mice fed diets containing unsterilized soil.

### Intake of soil microbes increased the abundance of genes for short-chain fatty acid metabolism and amino acid biosynthesis

Compared with that of the Control mice, the MW mice exhibited increased abundance of enzyme genes used in the metabolism of short-chain fatty acid (Fig. S4A and Table S9), including butyric acid, propionic acid, and acetic acid, as well as genes involved in amino acid biosynthesis. Compared with that of the SS mice, the Soil mice exhibited similar differences (Fig. S4B and Table S9).

The Soil mice ingested the same soil as the SS mice, but the soil-based microbes remained for the Soil group. Compared with that of the Control group, the Soil group of mice demonstrated increased abundance of not only genes for T3SS and two-component systems but also that for short-chain fatty acid metabolism and amino acid synthesis (Fig. S3). Furthermore, the abundance of genes for flagellar assembly (Fig. S5) and bacterial chemotaxis (Fig. S6), as well as additional genes for short-chain fatty acid metabolism (Fig. S7) and amino acid biosynthesis, was also increased in the Soil mice (Fig. S3).

The intake of soil microbes played an important role in increasing the abundance of genes involved in short-chain fatty acid metabolism and amino acid biosynthesis. Besides the common functions induced by the intake of either soil microbes or sterile soil, the intake of soil containing the microbes prompted the germination a greater number of different functional genes in addition to their increased relative abundance.

### Soil-intake or sterilized soil-intake decreased serum IgE levels

To analyze the impact of ingesting soil, sterilized soil, and soil microbe-containing water on the immune function of mice, we stimulated eczema on the skin of the four experimental groups of mice using DNFB and then measured serum IgE levels. The results revealed the serum IgE levels of the Soil and SS mice were significantly lower than those of the Control mice (*P* < 0.05). Furthermore, the levels of the Soil mice were significantly lower than the MW mice (*P* < 0.05; Fig. 5A). Although the median IgE value of the SS group was lower than the MW group, the difference was not statistically significant (Fig. 5A). Skin damage was also scored for the mice. The skin lesion scores in the Soil and SS groups were significantly lower than those in the Control and MW groups (*P* < 0.05), but there was no significant difference between the Soil and SS mice or between the MW and Control mice (Fig. 5B).

**Fig. 5.**
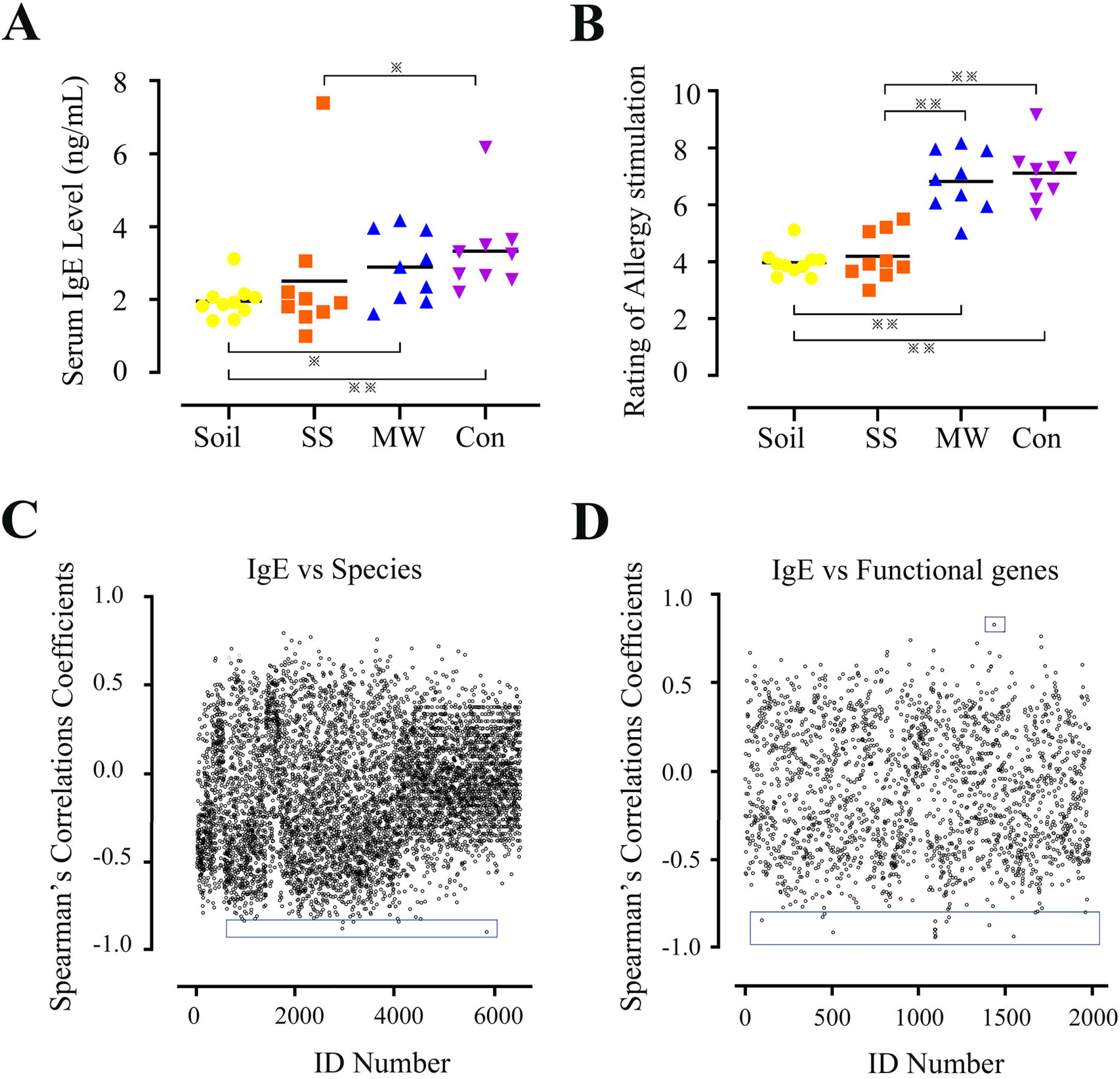
Effects of intake of soil, sterilized soil, or soil microbes on immunity by changing microbial community of gut. (A) Total serum IgE levels and (B) skin damage in mice treated with 2-4-dinitrofluorobenzene (DNFB) (n = 9 or 10/group; \**P* < 0.05, \*\**P* < 0.01, according to two-tailed least significant difference test). (C) Spearman correlations between serum IgE levels and the abundance of species and (D) between serum IgE levels and the abundance of function genes. The coefficients were calculated for the representation of each species or each function gene obtained from shotgun sequencing of the gut microbial metagenome. A spearman correlations coefficient of ± 1 indicates maximum correlation with age; zero indicates minimum correlation. The X-axis is the ID number of the species or the functional gene and the Y-axis is the correlation coefficient. Five microbes and seventeen genes with significant difference and their correlation coefficient are shown in square frame. Spearman correlation coefficients and P-values for all the species can be found in Table S11 and S12.

The Soil group and SS group demonstrated significant increases in the numbers of mouse pathogens and genes of T3SS. To determine whether these pathogens led to infection, hematological analysis of blood samples was performed. The results showed no indication of infection (Table S10).

To further explore which strain or gene was related to the enhancement of immune function in then mice, we performed a correlation analysis between IgE levels and the strains or functional genes detected by high-throughput sequencing (Fig. 5C, 5D, Table S11, and Table S12). The results showed that the microbes with significant correlation included fritillary virus Y and the soil bacteria *Burkholderia glumae*, among others (Table S11). The significance of these microbes in mice has not been reported, nor have there been any reports on their influence on immunity. The significant correlation for functional genes included six genes of T3SS and six genes of metabolic pathways (Table S12). The T3SS genes came from the same cell organ and work together to perform their function in promoting infection by the bacteria. Therefore, these T3SS-coding genes may be more closely related to the levels of IgE than the genes for the metabolic pathways.

There were 76 genes of bacterial secretion systems registered in the Kyoto Encyclopedia of Genes and Genomes (KEGG) database of which 43 (56.6%) were sequenced in this study. As shown in Table S13, 15 (34.9%) of the genes in the Soil group and 18 (41.9%) of the genes in the SS group were significantly more abundant than those in the Control group (*P* < 0.05). Only three genes of the MW group were observed at abundance higher than the Control group. Compared with that of the MW group, 11 (25.6%) genes of the Soil group and 9 (20.9%) genes of the SS group were more abundant.

Overall, the results showed that the intake of soil or sterilized soil improved the immune function of mice and did not cause obvious infections. The immune function was positively and correlated with the bacterial secretion system genes, especially with that of T3SS.

## Discussion

To explore the specific factors of the farm environment that influenced the protection against allergic diseases, we fed mice farm soil, sterilized soil, and soil microbes. The results showed that ingestion of sterilized soil significantly increased the diversity of intestinal microbes and abundance of T3SS genes, and decreased serum IgE levels, induced by DNFB. Ingestion of soil microbes increased the abundance of genes for the metabolism of short-chain fatty acids and the biosynthesis of amino acids. The intake of soil, which included the components of both the sterilized soil and the soil microbes, not only increased abundance in the intestinal microbiota of T3SS genes and the related infectious elements, but it also significantly increased the abundance of genes related to the metabolism of short-chain fatty acids and biosynthesis of amino acids. The intake of soil or sterilized soil significantly improved the immune function of mice and did not cause obvious infections. The immune function was positively and significantly correlated with the bacterial secretion system genes, especially with T3SS.

Based on our findings, soil played an important role in supporting the survival of some intestinal microorganisms in mice. In the absence of soil, ingestion of only soil microbes feeding resulted in changes of gut microbial diversity and structure, but the levels of change were significantly lower than those observed in the mice that ingested soil or sterilized soil. Therefore, soil may function as a “prebiotic” for various microbial strains or may be necessary, establishing a physical environment required for their survival. There are different ways soil functions: (1) soil is a mineral used for the survival of many intestinal microorganisms. Soil is known to contain abundant microorganisms (Lennon and Jones, 2011), at numbers 10 times greater than those in the human intestine (Lennon and Jones, 2011). (2) As a multi-pore structure, soil might provide the physical environment for microbial survival. Some microorganisms hide in soil particles, which may help them avoid been killed by microbial products such as antibiotics in the intestinal tract, or bactericides secreted by the human body such as antimicrobial peptides or IgA.

Ingestion of sterilized soil increased the diversity of intestinal microflora in mice under conditions in which the mice were inoculated with microorganisms via the air. Our previous study found that in a SPF animal facility, adding sterilized soil to mouse bedding changed the composition of intestinal microflora of mice, but did not increase the diversity of the intestinal microorganisms (Zhou et al., 2018). In the current study, the experiment was designed so the SS group of mice were reared next to the MW mice, which consumed soil microbes via their water. The open mouse cages could not prevent soil microbes from spreading to the SS mice. The results demonstrated increased gut microbial diversity of the SS mice.

In the current study, there was no significant difference in the immune function of mice fed soil microbes compared with that of the Control mice. Further, the intake of soil microbes significantly increased the abundance of genes involved in short-chain fatty acid metabolism and amino acid biosynthesis. Short-chain fatty acids play a role in improving human immunity (Furusawa et al., 2013). In addition, several reports have indicated that both resident and passing microbes can increase the immune function of mice (Rook, 2013; Honda and Littman, 2016). During ingesting soil microbes, many bacteria pass through the gut, which theoretically should stimulate the immune system of the mice and increase their immune function. There may be two reasons to explain our current results: (1) the Control group and MW group were raised in the same SPF animal facility. The mouse cage cover was a reticular structure and the soil microbes have spread from the cages of the MW mice to the Control mice, resulting in the immune function of the Control mice being enhanced; (2) we only evaluated the serum IgE levels for the mice after immune stimulation with DNFB and additional immune changes may not have been detected.

Pathogenic microbes and T3SS genes were detected in both the SS group and Soil group of mice, and the flagellar assembly gene was abundant in the Soil mice (Table S9 and Fig. S6). However, no infection was detected in the blood of any animals tested, which may have been due to these reasons: (1) the infection occurred during the early stage of the experiment and had resolved by the time we tested the blood of the mice; (2) the infection was weak and failed to present a clinical phenotype. No infections or deaths were observed during the experiment. It is very possible that the mice maintained a good interaction with the soil during the evolutionary process, or there may simply have been no strongly virulent pathogenic bacteria in the selected soil.

The current results also showed a strong correlation between IgE levels and the abundance of T3SS genes. T3SS exists in a variety of animal and plant pathogens, including Chlamydia spp. T3SS helps pathogenic microbes establish contact with host cells and plays roles in remodeling host cytoskeleton(Wong et al., 2012), host immune responses(Royan et al., 2010), and intestinal mucosal immunity(Kim et al., 2009). In addition, T3SS also exists in plant rhizobia and plays a key role in establishing symbiotic relationships with host cells(Büttner, 2012). The pathogenic mechanism of T3SS has been widely reported(Büttner, 2012), but the mechanism involved in improving host immune function needs further study.

In this study, a mouse model was used as the research approach, but there are great differences in lifestyles and evolutionary relationships between humans and mice. Therefore, the impact of soil on human intestinal microflora may differ and needs more experimental proof.

In conclusion, our results showed that an important reason farm environments have protective effects on allergic diseases is that soil can be used as a “prebiotic” to increase the diversity and richness of intestinal microbiota and promote the reproduction and growth of microorganisms harboring the genes of bacterial secretion system. Further mechanistic studies revealed that soil improved the natural immune function of mice mainly by increasing the abundance of genes of bacterial secretion system of gut microbiota, especially those of T3SS.

## Experimental Procedures

### Animals and experimental groups

At total of 60 mice aged 3–4 weeks were randomly divided into four groups (n = 20/group). The temperature of SPF animal facility was maintained at 24 ± 2 °C, humidity was 40 ± 5%, and the lights were on a 12 h/12 h light/dark cycle. Bedding material were change once a week. Starting at 7 weeks of age, the SS group was fed a diet containing 5% sterilized soil (Fig. S1), the Soil group was provided a diet containing 5% non-sterilized soil, and the MW group was provided drinking water containing ~10^11^ soil microbes. No treatment was performed for the Control group, which continued to receive a standard lab diet and normal drinking water.

After 42 d of treatment, feces from 10 mice in each group was randomly collected and stored at −80 °C for further intestinal microbiota analysis. Some mice in each group were used to perform the hematology analysis of blood samples, and the other mice were treated with DNFB and evaluated for serum IgE levels.

Soil was collected from farm ground at a depth greater than 0.5 cm, but no more than 10 cm. The soil composition was analyzed using a Wavelength Dispersive X-Ray Fluorescence spectrometer (Thermo Fisher, Waltham, MA, USA) with the results in Table S3. The soil was sterilized using autoclave at 121 °C for 30 min, which was repeated three times at a 24 h interval. Before use, the sterilized soil and non-sterilized soil were crushed and mixed with the mouse diet. Sterilized or non-sterilized soil feed was stored at −20 °C.

Soil microbes were isolated as following: Fresh farm soil was collected and mixed with sterile water at 2:1 (w/v). After stirring with a magnetic rod for 20 min, the solution was allowed to stand undisturbed for 10 min. The supernatant was then collected and centrifuged at 41 × *g* for 5 min. The supernatant was collected and allowed to stand for 48 h. The supernatant was again collected and centrifuged at 7440 × *g*. The supernatant was discarded, and the precipitate suspended in sterile water. A bacterial smear was prepared for microscopic examination. After confirming no soil particles remained in the microbe solution, it was added to the drinking water of the MW mice. The water was changed using a fresh microbial mixture once a week. The soil sample and microbes sample isolated from the soil underwent 16S rDNA high-throughput sequencing to analyze their microbial compositions (Table S2).

### Animal care and use

Animal experiments were conducted in strict accordance with the guidelines of the Animal Research Ethics Committee of Southeast University. All animal experiments were approved by the Animal Care Research Advisory Committee of Southeast University and the National Institute of Biological Sciences (approval number: 2017063009). All efforts were made to minimize and alleviate the pain the animals may experience. Specifically, the health of the mice was monitored every other day, and the weight measured weekly. The health status of mice was evaluated by observing changes in body weight and stool shape or appearance. In cases of serious diseases including one or more symptoms, such as weight loss of 15–20%, diarrhea, loss of hair quality, pain as indicated by an arched back or curled posture, or lethargy for over 1 week, the animals would be euthanized to reduce pain and suffering. Euthanasia was carried out: 1% (w/v) pentobarbital sodium (50 mg/kg) was injected intraperitoneally. Once the mice had lost consciousness, they were killed by a cervical dislocation and confirmed to be dead. Two people administered the injection, one held the animal, and the other gave the injection. The animals were not left unattended during the operation.

### Ethics approval

Animal experiments were carried out in strict accordance with the guidelines of the Animal Research Ethics Committee of Southeast University. All animal experiments were approved by the Animal Care Research Advisory Committee of Southeast University and the National Institute of Biological Sciences (approval number: 2017063009).

### DNFB treatment and serum IgE detection

DNFB was dissolved in acetone/olive oil (3:1) with the final concentrations of DNFB being 0.15% and 0.20%. Approximately 10 cm^2^ of hair was removed in the backs of the mice. After hair removal, 25 μl of 0.15% DNFB was applied to the ears of mice and 100 μl of the same solution was applied to the back on day 1 and day 4 of treatment. On days 7, 10, and 13, 0.2% DNFB was used to topically treat the mice at the same locations. Anesthesia of pentobarbital sodium was administered 24 h after the final DNFB treatment. Blood samples were collected from the eyes of the mice, allowed to clot, and the serum isolated through centrifugation. The serum was stored at −80 °C until analysis. Serum IgE levels were detected using an Invitrogen Mouse

IgE ELISA Ready-SET-Go Kit (eBioscience) following the manufacturer’s instructions.

### MiSeq sequencing and data handling

DNA sequencing and analysis were performed at Shanghai Biozeron Biological Technology Co. A fecal DNA Kit (Omega Bio-tek, Norcross, Georgia, USA) was used to extract the microbial DNA. The 515F and 907R primer set was used to PCR amplify the V4 hypervariable region of bacterial 16S rDNA and the amplicons were sequenced using an Illumina HiSeq X instrument with pair-end 150 bp (PE150) mode. FLASH software version 1.2.7 was used to merge the raw paired-end reads. Trimmomatic version 0.30 was used to eliminate low quality sequences. USEARCH version 7.1 was used to delete chimeric sequences. QIIME software version 1.9.1 software package was used to cluster the sequences into *de novo* OTUs using a 97% similarity threshold. RDP Classifier software version 11.5 was used to taxonomically assign the OTUs.

The genomic DNA was sheared using a Covaris S220 Focused-ultrasonicator (Woburn, MA USA). The fragmented DNA was then used to prepare metagenomic shotgun sequencing libraries. Trimmomatic was used to trim quality of the paw sequence reads and the Burrows-Wheeler Aligner mem algorithm used to map the sequences against the human genome. Clean taxonomic reads were then determined using Kraken2 according to the kraken database, which includes the National Center for Biotechnology Information (NCBI) RefSeq database in which all bacterial, archaeal, fungal, viral, protozoan, and algae genomic sequences are deposited (issue number: 90). The abundances of taxonomy were estimated using the Bayesian Reestimation of Abundance after Classification with KrakEN (Bracken) statistical method.

Megahit was used to generate a set of contigs for each sample. Prodigal (v2.6.3) was used to predict the open reading frames of the contigs. All open reading frames generated a set of unique genes after clustering with CD-HIT. The longest sequence of each cluster was the representative sequence for the genes. Salmon software was used to obtain the read number of each gene for calculating the gene abundance in the total sample. BLASTX was used to search the function of the genes coding proteins in the KEGG database. The specific functions and pathways were found in the KEGG path database.

### Data analysis and statistical tests

The α diversity indices were calculated for each sample based on the 16S rDNA sequence data using Chao1 estimator and the Shannon diversity/richness index. The 16S rDNA sequence data was also used to determine the KrakEN metrics of unweighted UniFrac distances between groups and PCoA was performed to show dissimilarities using QIIME. The Bray-Curtis dissimilarity metrics between any two groups was calculated based on the metagenomic sequence data and PCoA was performed using R version 3.2.3. Random forests analysis was performed as described previously (Yatsunenko et al., 2012). Briefly, R was used with 500 trees and all default settings to analyze the metagenomic sequence data of the 16S rDNA OTUs and species. Out-of-box error was used to estimate the generalization error. To calculate out-of-box error and importance score for each comparison, 10 relevant subsets of samples were extracted from OTU/species tables. SPSS software 18.0 (SPSS Company, Chicago, IL, USA) was used for t-test and analysis of variance (ANOVA).

## Data availability

Raw sequence reads for all samples described above were deposited into the NCBI Sequence Read Archive (SRA) database (project number, PRJNA686840)

## Acknowledgments

This work was supported by The Natural Science Foundation of China (grant no. 31770540), The Key Research Program of Jiangsu (grants no. BE2018663) and the Fundamental Research Funds for the Central Universities (grants no. 2242021k30014 and 2242021k30059).

## Conflict of Interest

No competing interests were disclosed.

## Notes

https://www.ncbi.nlm.nih.gov/sra/PRJNA686840

### Competing Interest Statement

The authors have declared no competing interest.

## References

A, P. (2003) Human geophagy: A review of its distribution, causes and implications. In: Skinner HCW, Berger AR, editors. Geology and Health: Closing the Gap. Oxford, UK: Oxford University Press.

Adlerberth, I., and Wold, A. (2009) Establishment of the gut microbiota in Western infants. Acta Paediatrica 98: 229–238.

Azad, M.B., Konya, T., Guttman, D.S., Field, C.J., Sears, M.R., HayGlass, K.T. et al. (2015) Infant gut microbiota and food sensitization: associations in the first year of life. Clin Exp Allergy 45: 632–643.

Büttner, D. (2012) Protein export according to schedule: architecture, assembly, and regulation of type III secretion systems from plant- and animal-pathogenic bacteria. Microbiol Mol Biol Rev 76: 262–310.

Berendsen, R.L., Pieterse, C.M., and Bakker, P.A. (2012) The rhizosphere microbiome and plant health. Trends Plant Sci 17: 478–486.

Blum, W.E.H., Zechmeister-Boltenstern, S., and Keiblinger, K.M. (2019) Does Soil Contribute to the Human Gut Microbiome? Microorganisms 7: 1–16.

Debarry, J., Garn, H., Hanuszkiewicz, A., Dickgreber, N., Blümer, N., von Mutius, E. et al. (2007) Acinetobacter Iwoffii and Lactococcus lactis strains isolated from farm cowsheds possess strong allergy-protective properties. J Allergy Clin Immunol 119: 1514–1521.

Ege, M.J., Mayer, M., Normand, A.-C., and Jon Genuneit, W.O.C.M.C., Charlotte Braun-Fahrländer, Dick Heederik, Renaud Piarroux, Erika von Mutius, GABRIELA Transregio 22 Study Group (2011) Exposure to Environmental Microorganisms and Childhood Asthma. New England Journal of Medicine 364: 701–709.

Fujimura, K.E., Johnson, C.C., Ownby, D.R., Cox, M.J., Brodie, E.L., Havstad, S.L. et al. (2010) Man’s best friend? The effect of pet ownership on house dust microbial communities. J Allergy Clin Immunol 126: 410–412, 412.e411-413.

Furusawa, Y., Obata, Y., Fukuda, S., Endo, T.A., Nakato, G., Takahashi, D. et al. (2013) Commensal microbe-derived butyrate induces the differentiation of colonic regulatory T cells. Nature 504: 446–450.

Gilbert, J.A., and Blaser, MJ. (2018) Current understanding of the human microbiome. 24: 392–400.

Grieneisen, L.E., Charpentier, M.J.E., Alberts, S.C., Blekhman, R., Bradburd, G., Tung, J., and Archie, E.A. (2019) Genes, geology and germs: Gut microbiota across a primate hybrid zone are explained by site soil properties, not host species. Proceedings of the Royal Society B: Biological ences 286.

Honda, K., and Littman, D.R. (2016) The microbiota in adaptive immune homeostasis and disease. Nature 535: 75–84.

Kembel, S.W., Jones, E., Kline, J., Northcutt, D., Stenson, J., Womack, A.M. et al. (2012) Architectural design influences the diversity and structure of the built environment microbiome. Isme j 6: 1469–1479.

Kim, M., Ogawa, M., Fujita, Y., Yoshikawa, Y., Nagai, T., Koyama, T. et al. (2009) Bacteria hijack integrin-?mked kinase to stabilize focal adhesions and block cell detachment. Nature 459: 578–582.

Lehtimäki, J., Thorsen, J., Rasmussen, M.A., Hjelmsø, M., Shah, S., Mortensen, M.S. et al. (2020) Urbanized microbiota in infants, immune constitution and later risk of atopic diseases. J Allergy Clin Immunol.

Lennon, J.T., and Jones, S.E. (2011) Microbial seed banks: the ecological and evolutionary implications of dormancy. Nat Rev Microbiol 9: 119–130.

Mhuireach, G., Johnson, B.R., Altrichter, A.E., Ladau, J., Meadow, J.F., Pollard, K.S., and Green, J.L. (2016) Urban greenness influences airborne bacterial community composition. Science of The Total Environment 571: 680–687.

Mutius, E.v., and Vercelli, D. (2010) Farm living: effects on childhood asthma and allergy. Nat Rev Immunol 10: 861–868.

Noora, O., Lasse, R., Alina, S., Hanna, S., Piia, K., Jenni, L.K. et al. (2018) Soil exposure modifies the gut microbiota and supports immune tolerance in a mouse model. Journal of Allergy & Clinical Immunology: S0091674918309345–.

Riedler, J., Braun-Fahrländer, C., Eder, W., Schreuer, M., Waser, M., Maisch, S. et al. (2001) Exposure to farming in early life and development of asthma and allergy: a cross-sectional survey. The Lancet 358: 1129–1133.

Rook, G.A. (2013) Regulation of the immune system by biodiversity from the natural environment: an ecosystem service essential to health. Proc Natl Acad Sci U S A 110: 18360–18367.

Royan, S.V., Jones, R.M., Koutsouris, A., Roxas, J.L., Falzari, K., Weflen, A.W. et al. (2010) Enteropathogenic E. coli non-LEE encoded effectors NleH1 and NleH2 attenuate NF-kB activation. Molecular microbiology 78: 1232–1245.

Schuijs, M.J., Willart, M.A., Vergote, K., Gras, D., Deswarte, K., Ege, M.J. et al. (2015) Farm dust and endotoxin protect against allergy through A20 induction in lung epithelial cells. Science 349: 1106–1110.

Sing, D., and Sing, C.F. (2010) Impact of direct soil exposures from airborne dust and geophagy on human health. International journal of environmental research and public health 7: 1205–1223.

Song, S.J., Lauber, C., Costello, E.K., Lozupone, C.A., Humphrey, G., Berg-Lyons, D. et al. (2013) Cohabiting family members share microbiota with one another and with their dogs. Elife 2:e00458.

Stephen-Victor, E., and Chatila, T.A. (2019) Regulation of oral immune tolerance by the microbiome in food allergy. Current Opinion in Immunology 60: 141–147.

Vatanen, T., Kostic, Aleksandar D., d’Hennezel, E., Siljander, H., Franzosa, Eric A., Yassour, M. et al. (2016) Variation in Microbiome LPS Immunogenicity Contributes to Autoimmunity in Humans. Cell 165: 842–853.

Vogel, K., Blümer, N., Korthals, M., Mittelstädt, J., Garn, H., Ege, M. et al. (2008) Animal shed Bacillus licheniformis spores possess allergy-protective as well as inflammatory properties. J Allergy Clin Immunol 122: 307–312, 312.e301-308.

von Hertzen, L., Hanski, I., and Haahtela, T. (2011) Natural immunity. Biodiversity loss and inflammatory diseases are two global megatrends that might be related. EMBO Rep 12: 1089–1093.

Wong, A.R., Raymond, B., Collins, J.W., Crepin, V.F., and Frankel, G. (2012) The enteropathogenic E. coli effector EspH promotes actin pedestal formation and elongation via WASP-interacting protein (WIP). Cell Microbiol 14: 1051–1070.

Wopereis, H., Oozeer, R., Knipping, K., Belzer, C., and Knol, J. (2014) The first thousand days - intestinal microbiology of early life: establishing a symbiosis. Pediatr Allergy Immunol 25: 428–438.

Yatsunenko, T., Rey, F.E., Manary, M.J., Trehan, L., Dominguez-Bello, M.G., Contreras, M. et al. (2012) Human gut microbiome viewed across age and geography. Nature 486: 222–227.

Zhou, D., Bai, Z., Zhang, H., Li, N., and Lu, Z. (2018) Soil is a key factor influencing gut microbiota and its effect is comparable to that exerted by diet for mice. F1000 Research 7: 1588.

Zhou, D., Zhang, H., Bai, Z., Zhang, A., Bai, F., Luo, X. et al. (2015) Exposure to soil, house dust and decaying plants increases gut microbial diversity and decreases serum immunoglobulin E levels in BALB/c mice. Environ Microbiology.

