## Supplemental Figure 1 for "Soil causes gut microbiota to flourish and total serum IgE levels to decrease in mice"

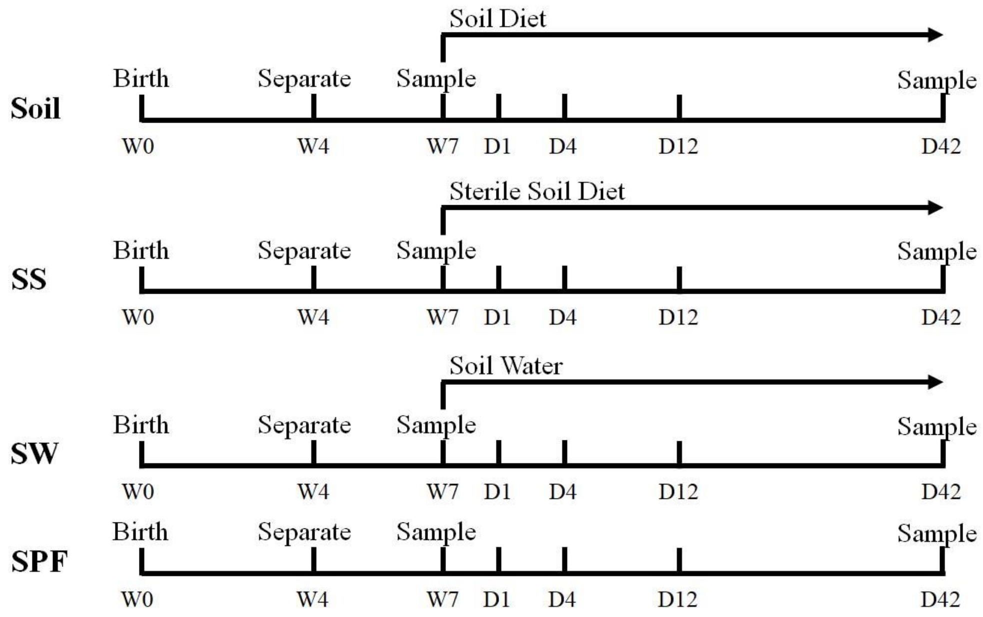


**Fig. S1.** Timeline indicating the treatment of mice with diets containing soil or sterile soil, or soil microbes provided in their drinking water, and the collection of fecal samples. For the three test groups, the treatments included feeding the mice diets containing soil (Soil) or sterile soil (SS), or providing soil microbes in the drinking water (MW). The treatments were administered starting at the age of 7 weeks (W7). Untreated animals served as a control group (Con). Fecal samples were collected on the 42nd day (D42) of treatments and the treatment started on day 0 (D0) for all the four groups of mice.
