## Supplemental Figure 2 for "Soil causes gut microbiota to flourish and total serum IgE levels to decrease in mice"

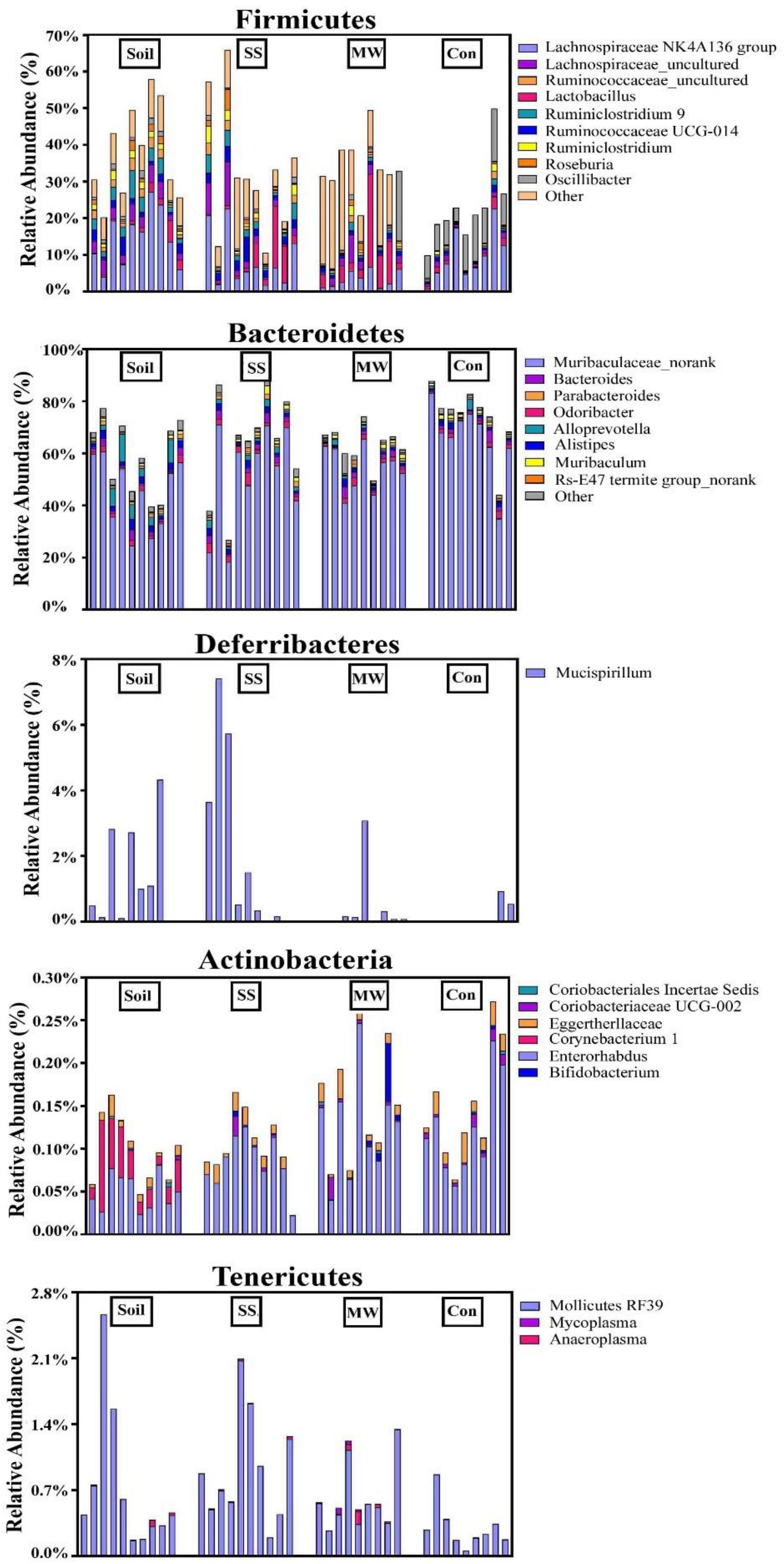


**Fig. S2.** Detailed relative abundance of bacterial genera classified via 16S rDNA sequences; the five most abundant major phyla of the gut microbiota observed:

A. Firmicutes. B. Bacteroidetes. C. Deferribacteres. D. Actinbacteria. E. Tenericutes. Each bar represents an individual mouse.
