## Supplemental Figure 3 for "Soil causes gut microbiota to flourish and total serum IgE levels to decrease in mice"

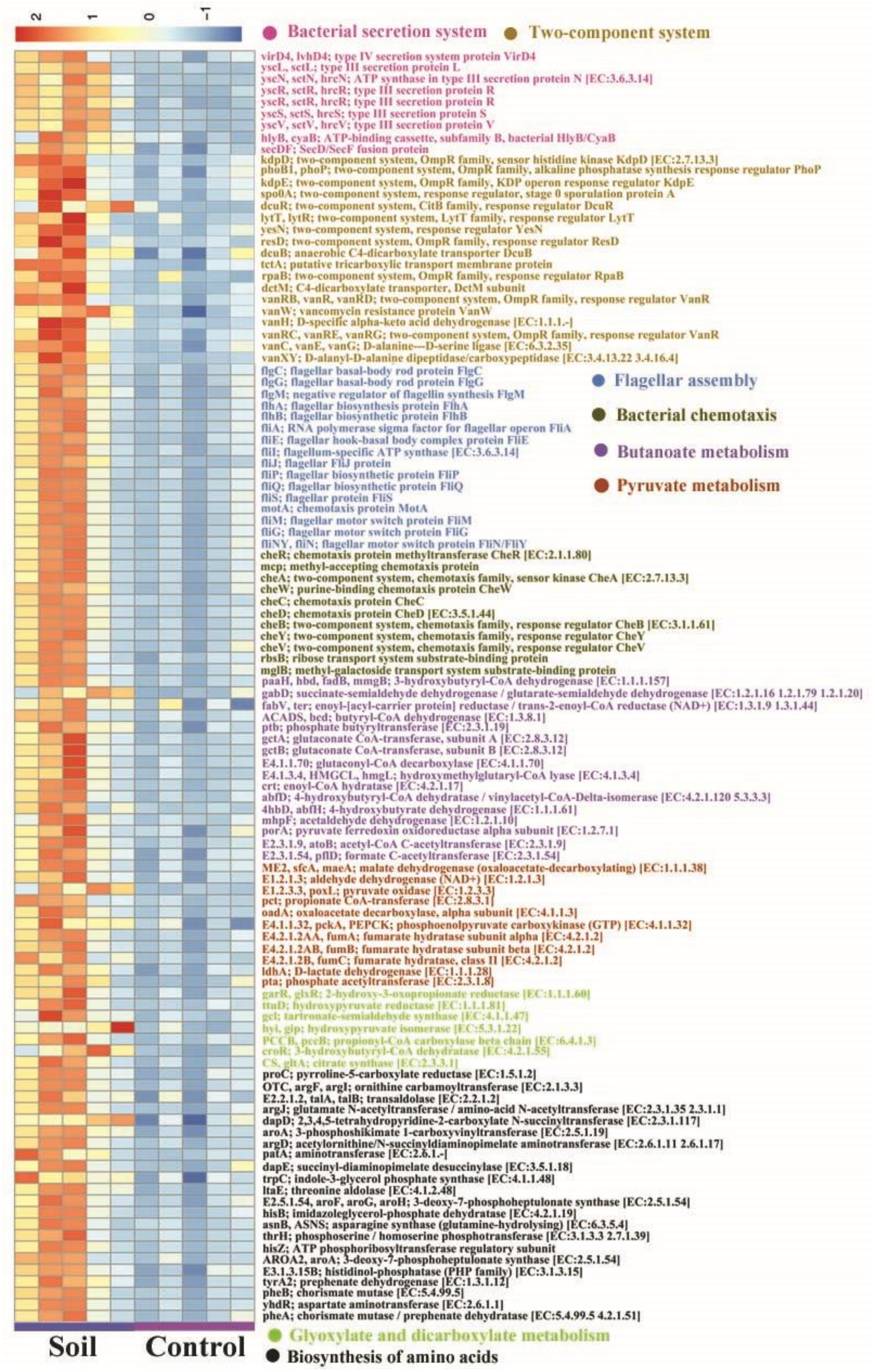


**Fig. S3.** Heatmap for abundant functional genes of intestinal microbiota of mice that ingested soil with their diet (Soil). The analysis was based on metagenomic shotgun sequencing data of the Soil group of mice compared to that of the Control group mice. (n = 5/group). The data are listed in Table S9.
