## Supplemental Figure 4 for "Soil causes gut microbiota to flourish and total serum IgE levels to decrease in mice"

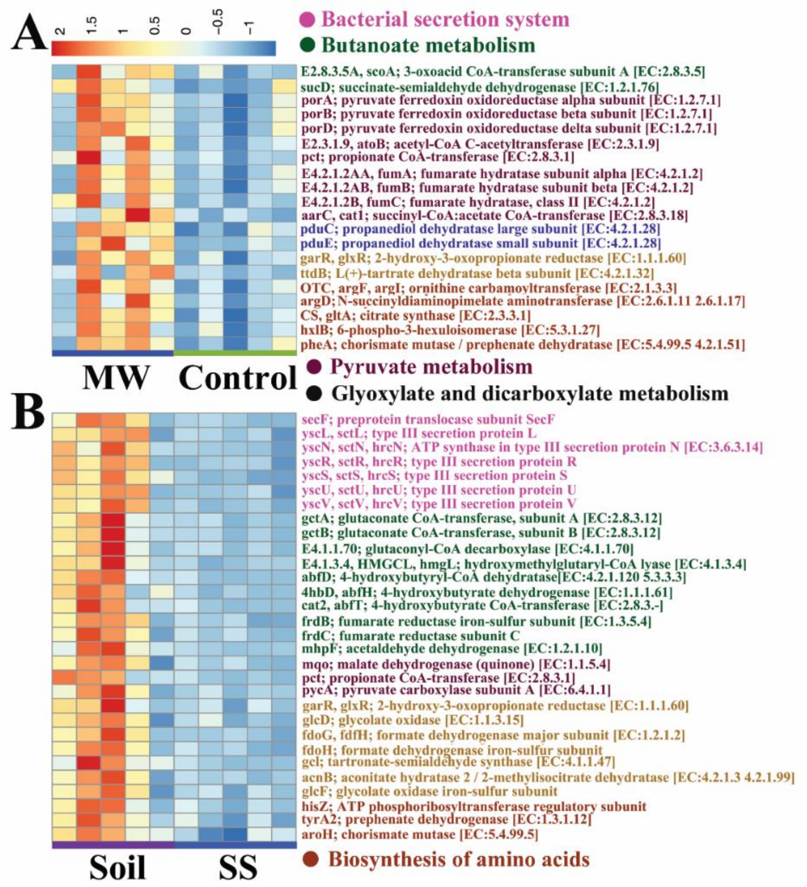


**Fig. S4. Heatmaps for effects of soil microbe-intake on gene abundance of gut microbial metagenomics**. A. Comparison between the MW and Control groups of mice (n = 5/group). B. Comparison between the Soil and SS groups of mice (n = 5/group). Treatment groups include mice receiving soil added to their diet (Soil) or sterile soil added to their diet (SS) or provided soil microbes in their drinking water (MW). Untreated mice served as a control (Control) group.
