## Supplemental Figure 7 for "Soil causes gut microbiota to flourish and total serum IgE levels to decrease in mice"

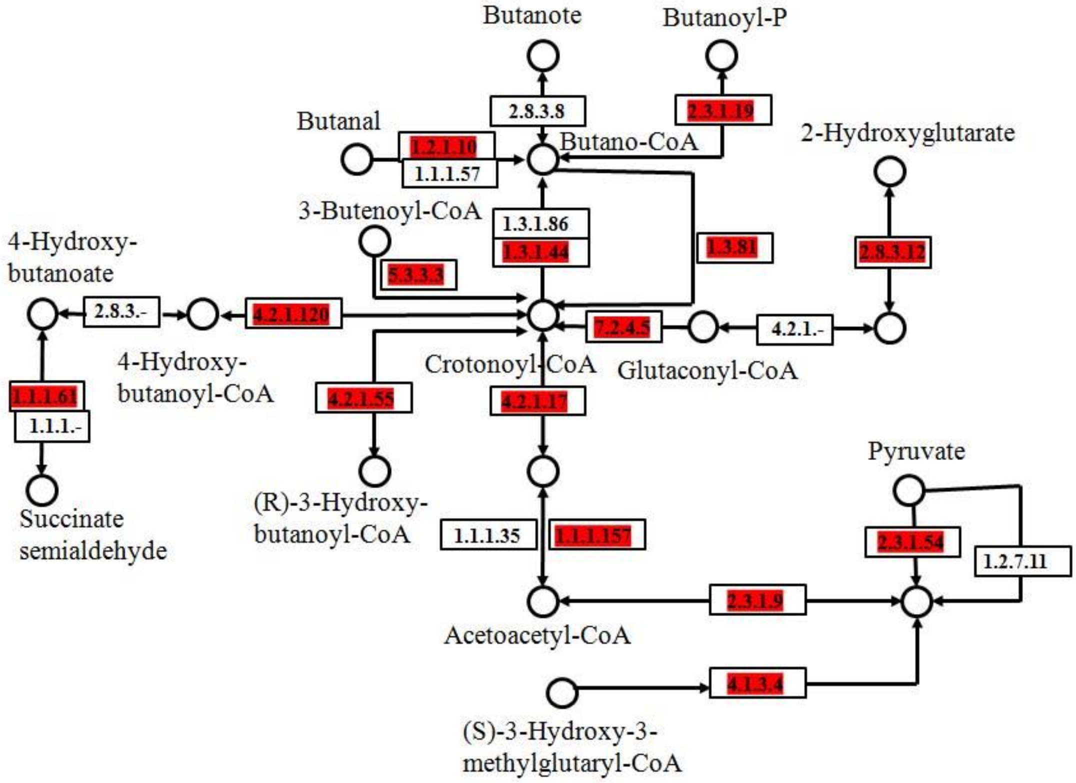


**Fig. S7.** Diagram of Kyoto Encyclopedia of Genes and Genomes (KEGG) pathway for butanoate metabolism. KEGG entries whose proportional representation was higher in the fecal microbiomes of the Soil mice compared with that in the Control mice. P-values for the highlighted KEGG entries can be found in Table S9. Control: experiment control mice; Soil: mice fed diets containing unsterilized soil.
