## Supplemental Tables for "Soil causes gut microbiota to flourish and total serum IgE levels to decrease in mice": ~WRL0510.tmp

**This file includes:**

Figures. S1 to S11

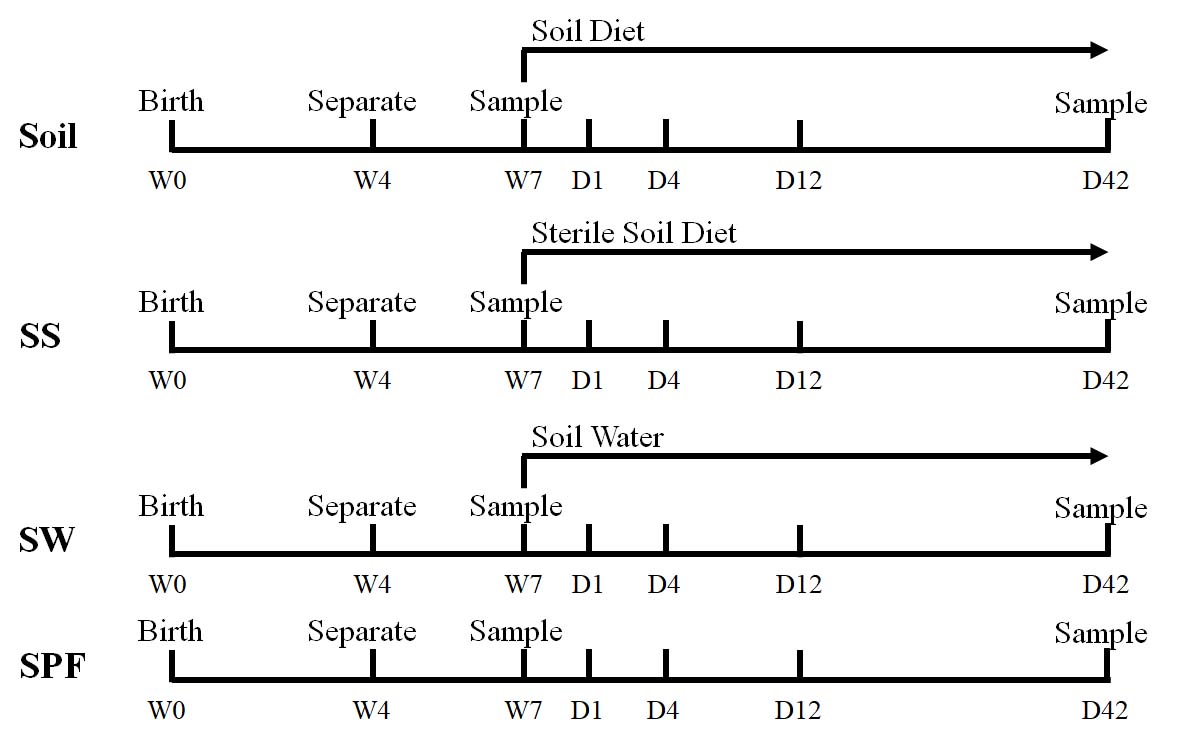

**Fig. S1.** Timeline indicating the treatment of mice with diets containing soil or sterile soil, or soil microbes provided in their drinking water, and the collection of fecal samples. For the three test groups, the treatments included feeding the mice diets containing soil (Soil) or sterile soil (SS), or providing soil microbes in the drinking water (MW). The treatments were administered starting at the age of 7 weeks (W7). Untreated animals served as a control group (Con). Fecal samples were collected on the 42nd day (D42) of treatments and the treatment started on day 0 (D0) for all the four groups of mice.

­­­­
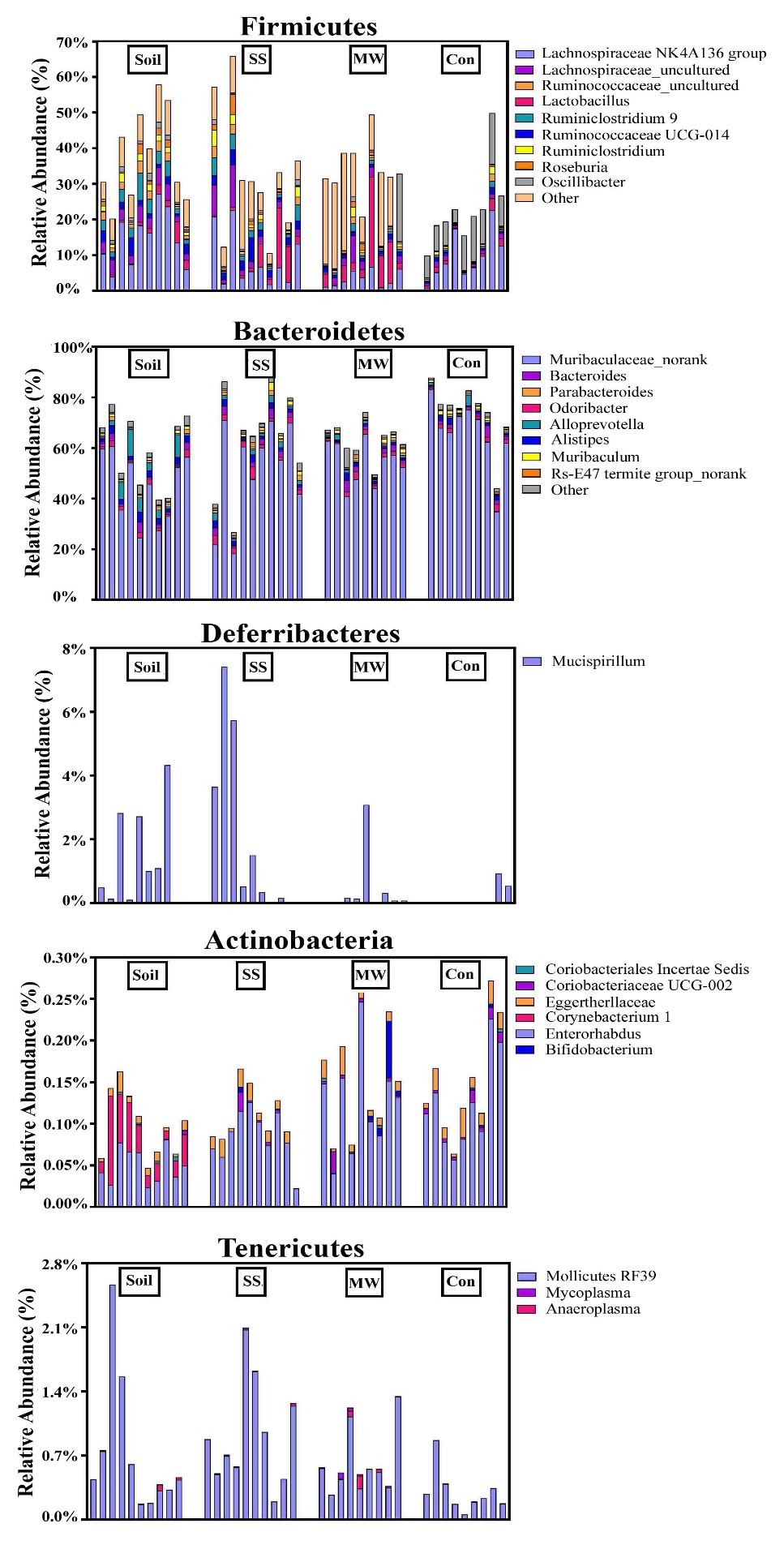

**Fig. S2.** Detailed relative abundance of bacterial genera classified via 16S rDNA sequences; the five most abundant major phyla of the gut microbiota observed:

A. Firmicutes. B. Bacteroidetes. C. Deferribacteres. D. Actinbacteria. E. Tenericutes. Each bar represents an individual mouse.

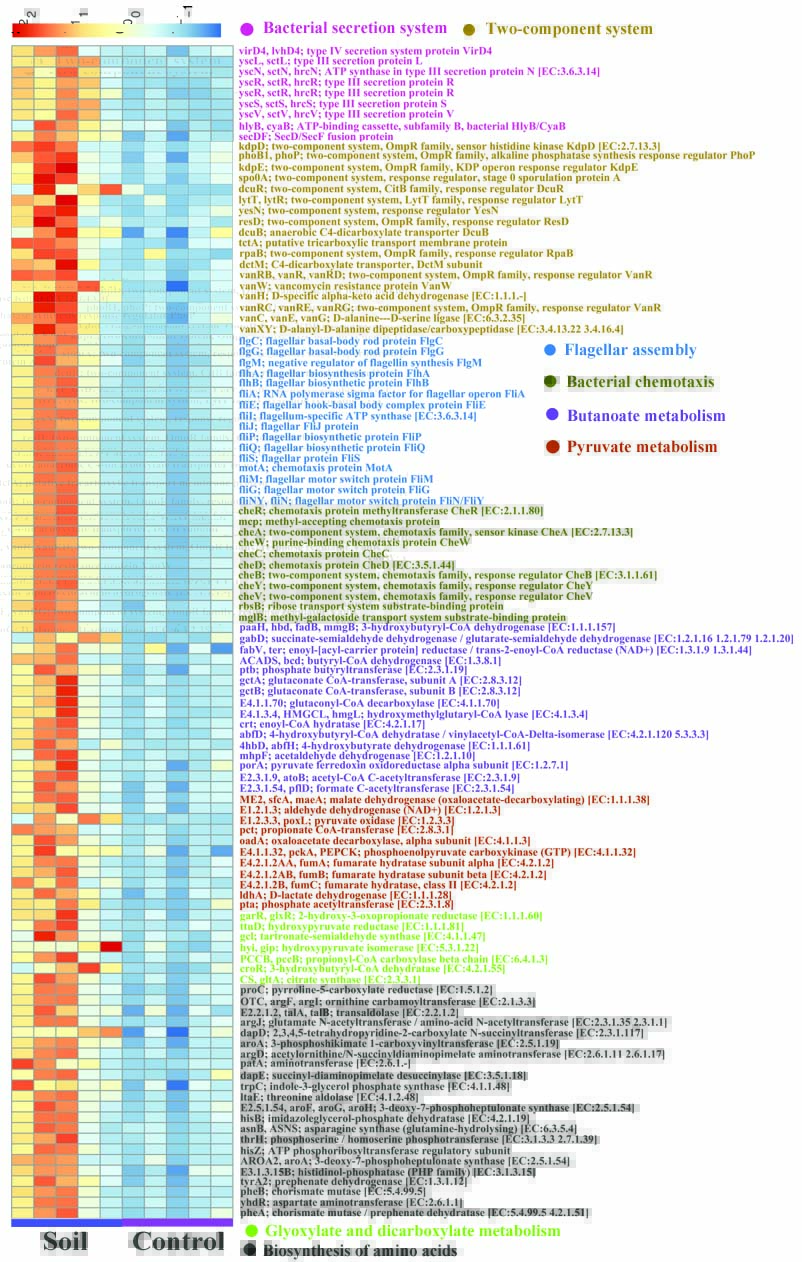

**Fig. S6.** Heatmap for abundant functional genes of intestinal microbiota of mice that ingested soil with their diet (Soil). The analysis was based on metagenomic shotgun sequencing data of the Soil group of mice compared to that of the Control group mice. (n = 5/group). The data are listed in Table S9.

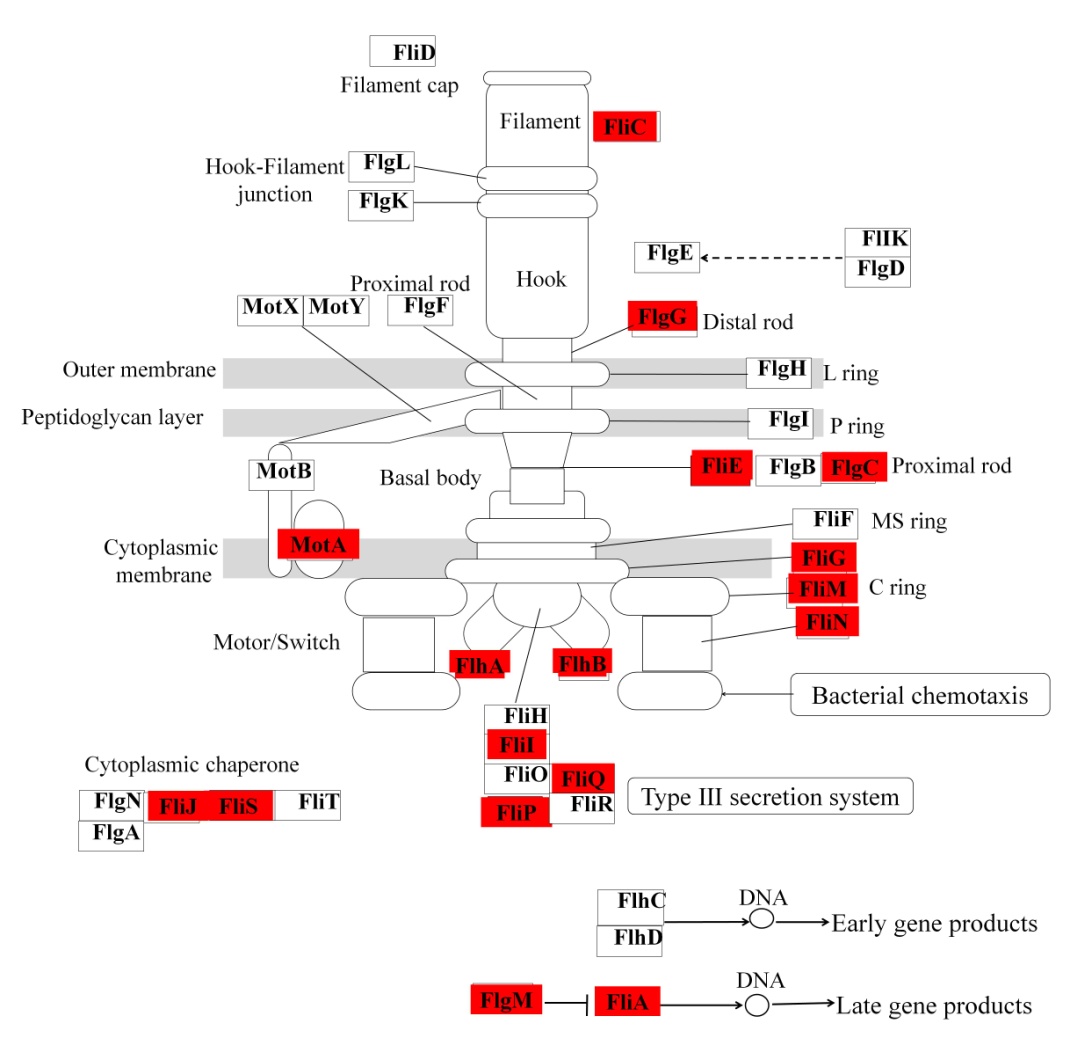

**Fig. S7.** Diagram of Kyoto Encyclopedia of Genes and Genomes (KEGG) entries for flagellar assembly indicating KEGG entries whose proportional representation was higher in the fecal microbiomes of the Soil group mice compared with that in the Control mice. P-values for the highlighted Kos can be found in Table S9. Control: experiment control mice; Soil: mice fed diets containing unsterilized soil.

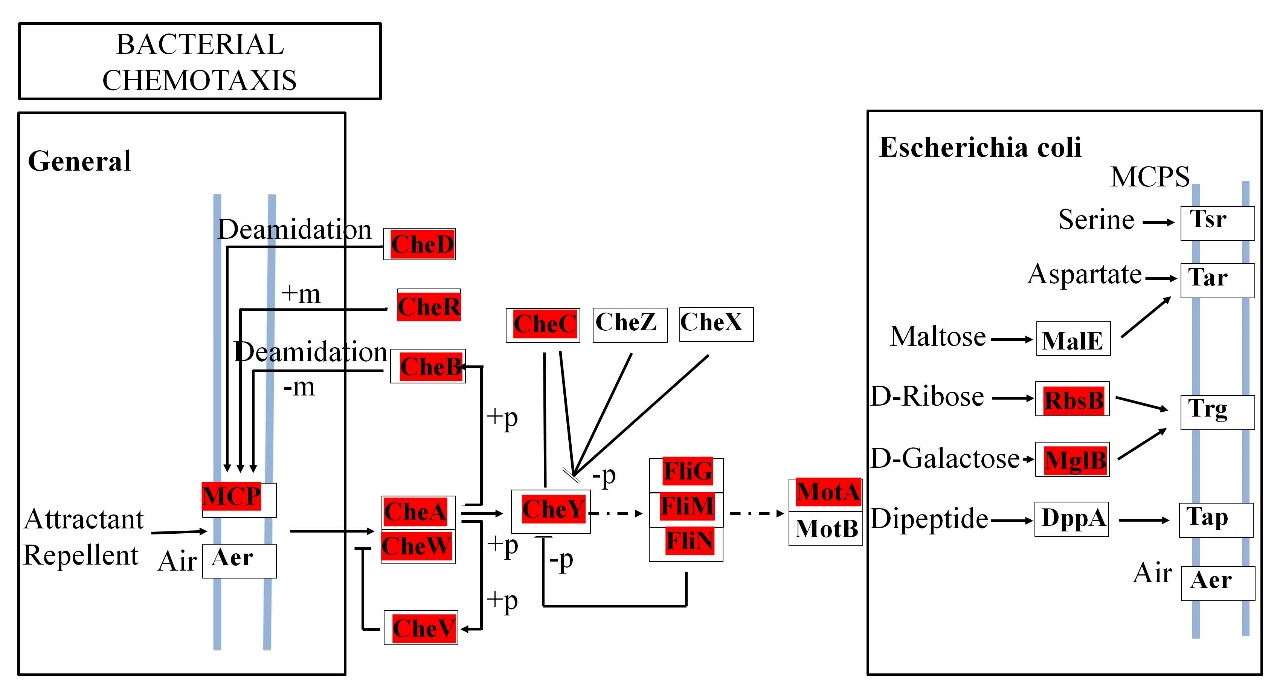

**Fig. S8.** Diagram of Kyoto Encyclopedia of Genes and Genomes (KEGG) entries for bacterial chemotaxis. KEGG entries whose proportional representation was higher in the fecal microbiomes of the Soil group mice compared with that in the Control mice. P-values for the highlighted KEGG entries can be found in Table S9. Control: experiment control mice; Soil: mice fed diets containing unsterilized soil.

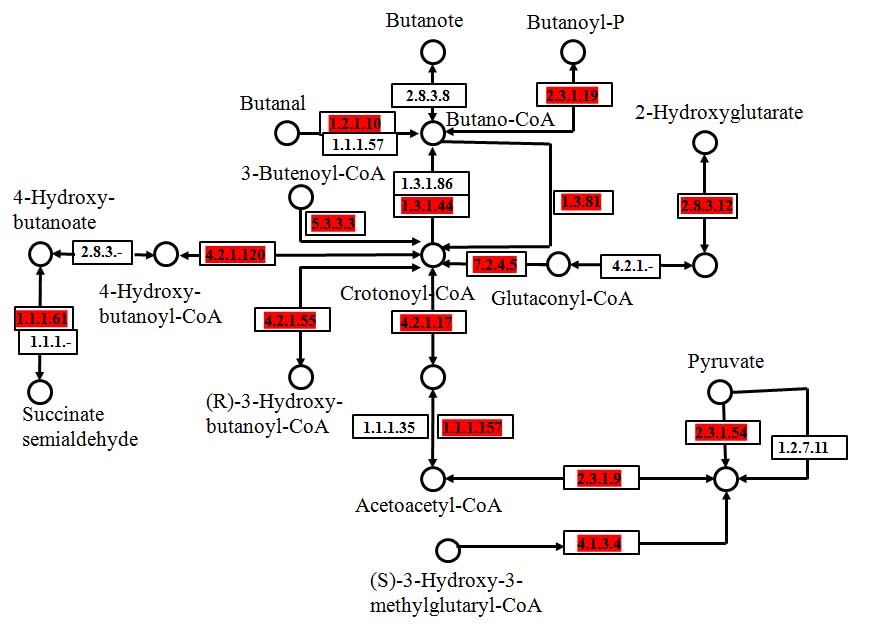

**Fig. S9.** Diagram of Kyoto Encyclopedia of Genes and Genomes (KEGG) pathway for butanoate metabolism. KEGG entries whose proportional representation was higher in the fecal microbiomes of the Soil mice compared with that in the Control mice. P-values for the highlighted KEGG entries can be found in Table S9. Control: experiment control mice; Soil: mice fed diets containing unsterilized soil.

**Fig. S10.** Spearman correlations between serum IgE levels and the abundance of species. The coefficients were calculated for the representation of each species obtained from shotgun sequencing of the gut microbial metagenome. A spearman correlations coefficient of ± 1 indicates maximum correlation with age; zero indicates minimum correlation. The X-axis is the ID number of the species and the Y-axis is the correlation coefficient. Five microbes with significant difference and their correlation coefficient being at least 0.88 are shown. Spearman correlations coefficients and P-values for all the species can be found in Table S11.

**Fig. S11.** Spearman correlations between serum IgE levels and the abundance of function genes. The coefficients were calculated for the representation of each function gene obtained from shotgun sequencing of the gut microbial metagenome. A spearman correlations coefficient of ± 1 indicates maximum correlation with age; zero indicates minimum correlation. The X-axis is the ID number of the functional gene and the Y-axis is the correlation coefficient. The function genes with spearman correlation coefficients under 0.80 have significant differences. Spearman correlation coefficients and P-values for all the species can be found in Table S11.
